# Assessing tACS-induced phosphene perception using closed-loop Bayesian optimization

**DOI:** 10.1101/150086

**Authors:** Romy Lorenz, Laura E. Simmons, Ricardo P. Monti, Joy L. Arthur, Severin Limal, Ilkka Laakso, Robert Leech, Ines Violante

**Affiliations:** Computational, Cognitive and Clinical Neuroscience Laboratory (C3NL), Department of Medicine, Imperial College London, London W12 0NN, UK; Department of Bioengineering, Imperial College London, London SW7 2AZ, UK; Gatsby Computational Neuroscience Unit, University College London, W1T 4JG London, UK; Department of Electrical Engineering and Automation, Aalto University, 02150 Espoo, Finland

## Abstract

Transcranial alternating current stimulation (tACS) can evoke illusory flash-like visual percepts known as *phosphenes*. The perception of phosphenes represents a major experimental challenge when studying tACS-induced effects on cognitive performance. Besides growing concerns that retinal phosphenes themselves could potentially have neuromodulatory effects, the perception of phosphenes may also modify the alertness of participants. Past research has shown that stimulation intensity, frequency and electrode montage affect phosphene perception. However, to date, the effect of an additional tACS parameter on phosphene perception has been completely overlooked: the relative phase difference between stimulation electrodes. This is a crucial and timely topic given the confounding nature of phosphene perception and the increasing number of studies reporting changes in cognitive function following tACS phase manipulations. However, studying phosphene perception for different frequencies and phases simultaneously is not tractable using standard approaches, as the physiologically plausible range of parameters results in a combinatorial explosion of experimental conditions, yielding impracticable experiment durations. To overcome this limitation, here we applied a Bayesian optimization approach to efficiently sample an exhaustive tACS parameter space. Moreover, unlike conventional methodology, which involves subjects judging the perceived phosphene intensity on a rating scale, our study leveraged the strength of human perception by having the optimization driven based on a subject’s relative judgement. Applying Bayesian optimization for two different montages, we found that phosphene perception was affected by differences in the relative phase between cortical electrodes. The results were replicated in a second study involving new participants and validated using computational modelling. In summary, our results have important implications for the experimental design and conclusions drawn from future tACS studies investigating the effects of phase on cognition.

## Introduction

Externally induced oscillations generated by transcranial alternating current stimulation (tACS) offer a way to directly manipulate rhythmic activity in the brain (Antal and Paulus, 2013; Herrmann et al., 2013). The rationale behind tACS is that with a continuous oscillatory stimulation, the spontaneous activity of an increasing number of individual neurons starts to synchronize with the external oscillator, a phenomenon known as *neural entrainment* (Ali et al., 2013; Fröhlich and McCormick, 2010; Helfrich et al., 2014). This property makes tACS a powerful technique for studying causal relationships between neural oscillations and behaviour (Antonenko et al., 2016; Polanía et al., 2012; Violante et al., 2017), while also holding promise for therapeutic uses (Brittain et al., 2013; Schmidt et al., 2013; Wu et al., 2016).

It is well established that tACS can evoke illusory flash-like visual percepts known as *phosphenes*. The current understanding is that phosphenes are generated in the retina by current spreading from the stimulation electrodes. Alongside empirical work (Kar and Krekelberg, 2012; Schutter, 2016; Schutter and Hortensius, 2010; Schwiedrzik, 2009), this hypothesis is supported by computational models suggesting that even distant electrodes can result in small portions of current flowing through the eyes, sufficient to induce retinal phosphenes due to the high sensitivity of the retina to electrical stimulation (Laakso and Hirata, 2013). However, alongside the evidence for the retinal origin of phosphene perceptions, possible interaction effects with ongoing cortical oscillations cannot be fully dismissed. For example, Terhune et al. (2015) reported lower tACS-induced phosphene thresholds at 40 Hz for a specific type of synaesthetes (i.e., “projectors”) in comparison to another type of synaesthetes (i.e., “associators”) or non-synaesthete controls. The authors concluded that observed differences in phosphene thresholds are likely to be of cortical origin, as neuroanatomical differences in the primary visual cortex have been found for “projector”-type synaesthetes.

The importance of studying phosphene perception induced by tACS was recently highlighted by Schutter (2016), who pointed to an emergent body of evidence suggesting that retinal phosphenes can induce neural entrainment. Therefore, visual phosphenes may be both a byproduct of entrainment and may themselves cause patterns of entrainment in the same way as exogenous photic stimulation (Schutter, 2016). Importantly, photic stimulation has been shown to improve cognitive performance (Elliott and Müller, 1998; Knez, 2014; Williams et al., 2006; Williams, 2001). These findings indicate a possible confounding factor of tACS-induced changes in behavioural and cognitive improvements, since they may not be “solely attributable to the direct neuromodulatory effects” (Schutter, 2016) of tACS, but could potentially be caused by retinal phosphenes.

To date, three tACS-related parameters have been shown to affect the perception of phosphenes: the frequency and intensity of stimulation as well as electrode montage. Phosphene perception is highly dependent on the stimulation frequency, with the strongest perception reported in the lower beta range, i.e., 14-22 Hz (Kanai et al., 2008; Turi et al., 2013). Equally, phosphene perception is more pronounced the higher the stimulation intensity (Cabral-Calderin et al., 2016; Kanai et al., 2008; Schutter and Hortensius, 2010). While it was found that phosphene perception greatly depends on the exact placement of active and return electrodes (Mehta et al., 2015), the consensus is that the closer the tACS electrodes are placed to the eye, the stronger the perception of phosphenes will be (Kar and Krekelberg, 2012; Laakso and Hirata, 2013; Paulus, 2011; Schutter and Hortensius, 2010); these observations are also in line with the hypothesis of a retinal origin of phosphenes.

However, there is an additional parameter with the potential to impact phosphene perception that has not yet been investigated: the effect of relative phase difference between independent electrodes. Relative phase refers to the difference in phase angle between oscillations, and can be simply understood as the time lag between oscillators. The relative phase between brain oscillations has been shown to affect perception and cognitive function (Antal and Herrmann, 2016). For example, tACS induced theta phase synchronization (i.e., coupling between oscillations at theta frequencies with ~0° relative phase difference) over frontoparietal regions was found to improve working memory performance (Polanía et al., 2012; Violante et al., 2017). Moreover, the relative phase of delta oscillations plays a role in speech recognition (Henry and Obleser, 2012) and gamma phase coupling between hemispheres has been associated with an enhanced perception of horizontal motion, which is necessary for efficient inter-hemispheric integration (Rose and Büchel, 2005).

A practical challenge when aiming to assess phosphene perception for different frequencies and phases simultaneously is that it rapidly results in a combinatorial explosion of experimental conditions, as physiologically plausible frequencies (0.1-100 Hz) and phases (0-359°) span across a wide range of possibilities; thus, resulting in impracticable experiment durations. Furthermore, the conventional approach for assessing phosphene perception typically has subjects judging perceived intensity on a rating scale^1^ (e.g., 0-3 (Kanai et al., 2008; Schutter and Hortensius, 2010), 1-4 (Pascual-Leone and Walsh, 2001; Silvanto and Cattaneo, 2010), 0-5 (Cabral-Calderin et al., 2016), or 0-10 (Mehta et al., 2015)). This is problematic as it relies on judgments about absolute magnitudes; however, humans are better at making relative judgments (Kahneman and Tversky, 1979; Miller, 1956; Seymour and McClure, 2008; Stewart et al., 2001, 2005), expressing preference for one option over another.

To overcome these limitations, here we applied Bayesian optimization (Lorenz et al., 2016b, 2017a) to efficiently search through an exhaustive tACS parameter space, with the aim of identifying frequency-phase combinations that elicited the strongest phosphene perception in individual subjects. Bayesian optimization is an active learning technique that is characterized by automatically choosing samples, from which it progressively learns in real-time. Active sampling is particularly useful for scenarios when we are interested in exploring a large space of possible experimental conditions but where the acquisition of appropriate training data comes at a cost, either in terms of time, financial costs, data quality and/or subject comfort. In case of behavioural experiments with human participants, sampling all possible tACS frequency-phase combinations would be practically unfeasible, as the total length of tACS stimulation would exceed the recommended session duration (Fregni et al., 2015). In addition, highly repetitive behavioural tasks quickly lead to subject disengagement, possibly impairing data quality. Bayesian optimization intelligently and efficiently searches through a larger space of experimental conditions than is feasible with standard approaches.

In our study, at each iteration, subjects were exposed to two blocks of tACS with different frequency-phase-combinations, and the optimization was driven based on the subject’s rating of which of two blocks evoked the strongest phosphene perception (block 1 vs. block 2). Based on the subject’s choice, the algorithm automatically proposed a new pair of stimulation parameters to be applied to the subject in the next iteration (Figure 1). The empirical data obtained were subsequently compared to the results of a new computational model predicting current density in the retinas based on anatomical information of electrode placement (Laakso and Hirata, 2013), as well as differences in relative phase.

**Figure 1.**
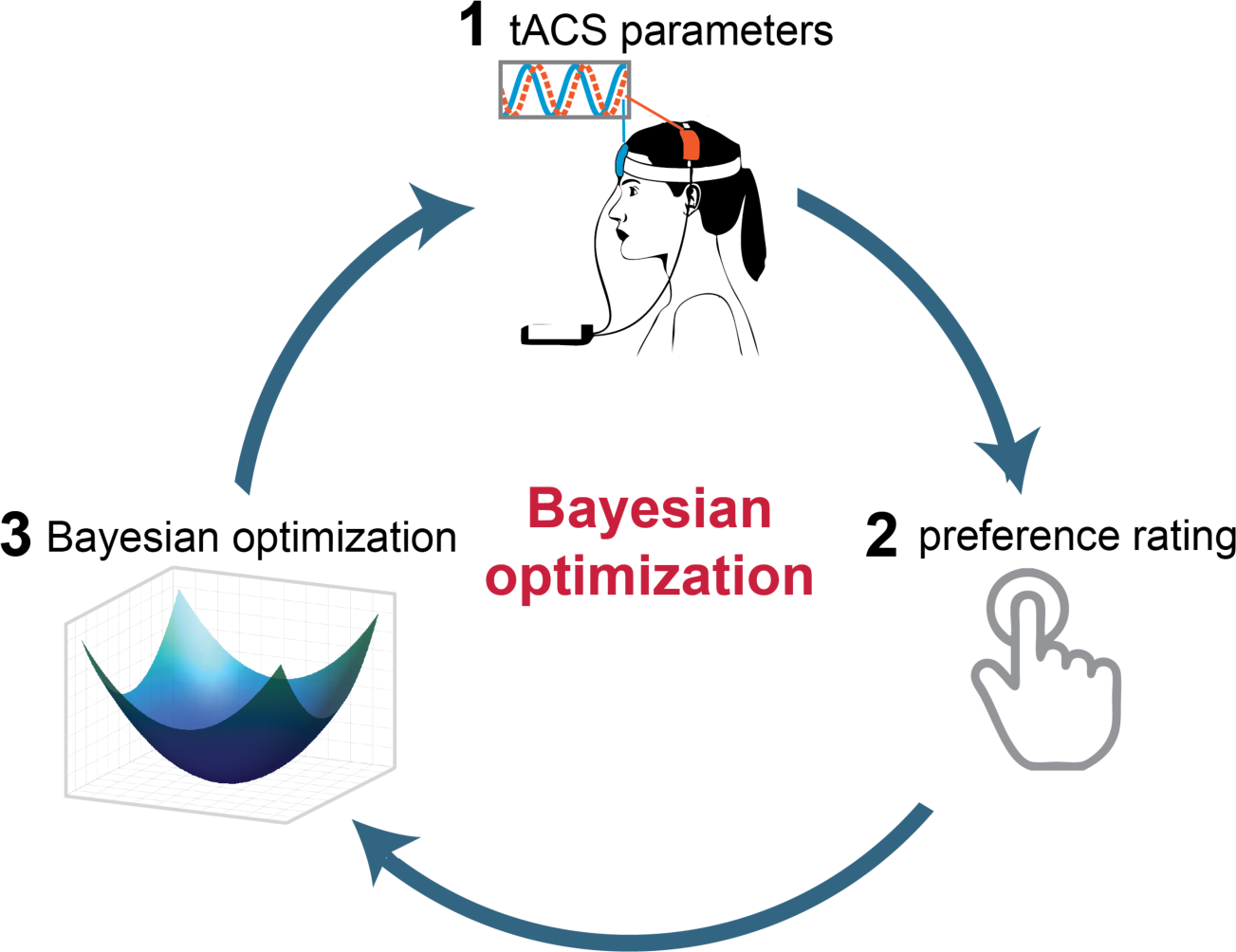
Bayesian optimization for assessing phosphene perception. (1) The experiment starts by the subject receiving two blocks of tACS; each block corresponds to a different tACS frequency-phase combination. (2) After both blocks, subjects indicate for which block the phosphene perception was stronger by pressing a button on a response box. (3) Based on the subject’s choice, the algorithm automatically proposes a new pair of stimulation parameters to be applied to the subject in the next iteration. This cycle continues until a stopping criterion is reached or the experiment is ended automatically.

## Methods

We performed two studies with the objective of investigating the phase-dependency of phosphene perception and demonstrating the feasibility of Bayesian optimization for assessing tACS-induced phosphene perception across a large tACS parameter space. In Study 1, three different montages were tested: midline central-occipital (Cz-Oz), right frontoparietal (F4-P4), and bilateral occipital (O1-O2). For montage Cz-Oz, perception of phosphenes was only tested across different frequencies. This was motivated by previous studies employing the same montage to assess the frequency-dependency of phosphene perception using conventional rating scales (e.g., (Kanai et al., 2008)). Therefore, the results obtained from the present study using a Bayesian optimization approach based on preference ratings could be directly compared to previous findings. For montages F4-P4 and O1-O2 phosphene perception was assessed for different tACS frequency-phase-combinations. The results of Study 1 served as a first validation step of the proposed method and were used to obtain more accurate priors for a second online study. For Study 2, new subjects were recruited and phosphene perception was assessed at a higher level of detail for montages F4-P4 and O1-O2 by using a smaller parameter space involving a narrower frequency-range. Study 2 served two purposes: 1) testing the replicability of results obtained on new data whilst; 2) having the optimization algorithm more thoroughly explore the phase-dependency of phosphene perception. Empirical data obtained were subsequently compared to a new computational model.

### Subjects

Twenty healthy volunteers took part in the two studies of ten participants each (Study 1: 6 females, mean age ± SD: 26.3 ± 5.59 years, range: 21-40 years, 1 left-handed and 1 ambidexter; Study 2: 5 females, mean age ± SD:26. 6 ± 4.69 years, range: 22-36 years, 2 left-handed). All participants had normal or corrected-to-normal vision, no history of epilepsy, traumatic brain injury or skull metal implants. Additionally, pregnant women and those with serious visual impairment were not asked to take part. Subjects gave written informed consent for their participation. The study conformed to the Declaration of Helsinki and ethical approval was granted through the local ethics board (NRES Committee London – West London & GTAC).

### TACS montage

Stimulation was delivered using two MR-compatible battery-driven stimulators (NeuroConn GmbH, Ilmenau, Germany) and controlled using Matlab via a data acquisition device (National Instruments, Newbury, UK). Stimulation was sinusoidal, with peak-to-peak amplitude of 1 mA, and no DC offset. For both studies, a within-subject design was applied, testing phosphene perception with different montages in separate runs for each participant. For the different montages, rubber electrodes (Ten20, D.O. Weaver, Aurora, CO, USA) were positioned on the scalp based on locations according to the International 10-20 EEG system (Klem et al., 1999). For Study 1, three different montages were tested (Figure 2a): Cz-Oz, F4-P4 and O1-O2. For the O1-O2 montage, return electrodes were placed on the left and right shoulders. For F4-P4, both return electrodes were placed ipsilaterally on the right shoulder. The Cz-Oz montage used only two electrodes, with the larger return electrode at the Cz position (for this montage stimulation was delivered using one stimulator). For Study 2, only two montages were considered: F4-P4, and O1-O2. The target stimulation electrodes were round (diameter = 4.3 cm, thickness: 1mm) while return electrodes were of a rectangular shape (7.3 x 5.4 cm, thickness: 2 mm). Impedances were kept below 10 kΩ using a conductive paste (Ten20, D.O. Weaver, Aurora, CO, USA), which also held the electrodes in place. A balanced Latin square design was used to alternate the order in which subjects were tested with each montage. Between montages, alcohol wipes were used to remove residue of the conductive paste from the scalp.

**Figure 2.**
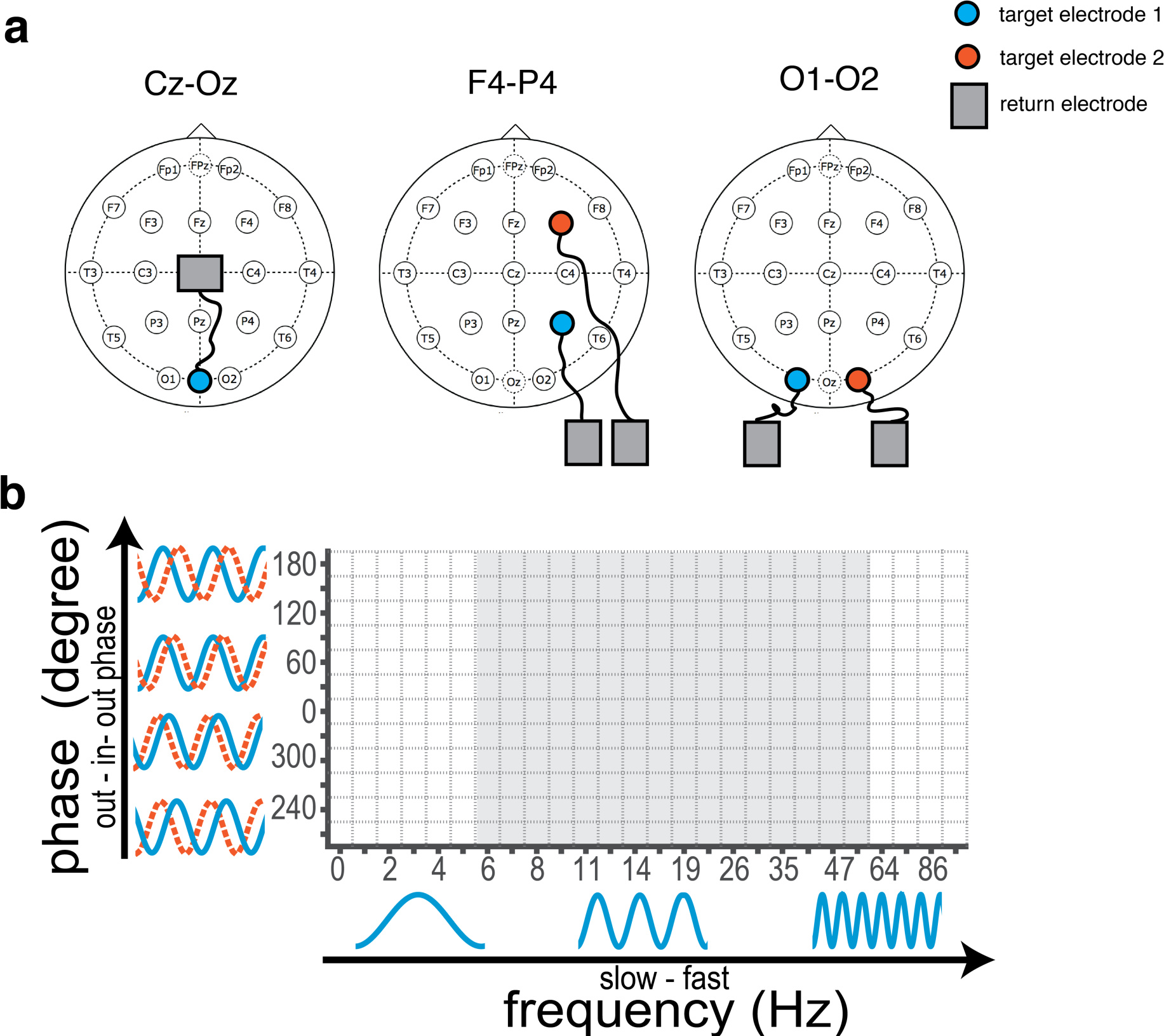
Experimental methodology. (a) In Study 1, three different montages were tested: Cz-Oz, F4-P4, and O1-O2. In Study 2, only the latter two were investigated. (b) In Study 1, the tACS parameter space searched by the Bayesian optimization algorithm consisted of 26x12 different tACS frequency-phase combinations (white and grey space). Study 2 zoomed into the parameter space by narrowing the frequency range considered, resulting in 16x12 different tACS frequency-phase combinations (grey space).

### TACS parameter space

In Study 1, montage Cz-Oz served to validate the technique against previous studies (e.g., (Kanai et al., 2008)) and was therefore optimized for a 1D parameter space (26x1) consisting of logarithmically spaced frequencies ranging from 0 to 100 Hz. For the two other montages, an exhaustive 2D parameter space (26x12) was designed, adding a further dimension consisting of relative phase differences ranging from 0-330° (Figure 2b). Based on the results of Study 1, Study 2 zoomed into the parameter space by narrowing the frequency range considered (6 – 55 Hz, grey shaded area in Figure 2b).

### Experimental Procedure

Subjects were seated centrally in front of a computer screen at a distance of 75 cm. The lights in the room were kept on throughout. For each of three electrode montages, subjects were habituated to the stimulation with the intensity increasing incrementally (0.25 mA, 0.5 mA and 1 mA). The first run was preceded by 5 s of stimulation at 16 Hz (0°); it has been reported that the majority of individuals will have phosphene perception at this frequency (Kanai et al., 2008), therefore this served to demonstrate to subjects what phosphene perception “looks” like.

Each run consisted of 20 rating blocks. At each iteration, subjects were exposed to two blocks of tACS lasting 5 s each, with a 1 s rest in between blocks, and a 5 s rest between iterations. During the stimulation, subjects were asked to fixate on the words “1.block” or “2.block” as they appeared on the screen (white font on grey background); during the rest period, they were asked to fixate on a white cross in the centre of the screen (grey background). After each pair of blocks, subjects were prompted to select the block in which they felt their perception of phosphenes had been stronger. The concept of “stronger” perception was explained before the experiment as: the phosphenes appearing brighter; the phosphenes taking over a larger area of the visual field; phosphenes appearing in both eyes where previously they had only appeared in one; or phosphenes appearing to be more intense or flickering more rapidly. Subjects made their choice between block 1 and 2 by pressing either the left or right button on a button response box. If subjects perceived no clear difference between rating blocks, or did not perceive any phosphenes, they were instructed to select a block at random. Based on the subject’s preference rating, the Bayesian optimization algorithm automatically proposed a new pair of stimulation parameters to be presented to the subjects in the next iteration. Each run lasted 5.5 minutes.

After each run, participants completed a questionnaire to assess possible side-effects of the tACS stimulation by rating from 0 (none) to 4 (severe) the intensity and duration of: pain, burning, warmth/heat, itchiness, pitching, metallic taste, fatigue, effect on performance or any other side-effect perceived. Additionally, subjects were asked to draw where in their visual field they perceived the phosphenes. The experimenter noted when subjects reported that they did not experience any phosphenes during the entire run. For montage Cz-Oz in Study 1, four subjects reported that they did not perceive any phosphenes over the course of the whole run. These subjects were excluded from subsequent analyses related to this specific montage. For the two other montages, all subjects reported phosphenes in both studies.

### Bayesian optimization for binary observations

Bayesian optimization can be understood as a two-stage procedure that repeats in an iterative closed-loop. The first stage is the *data modelling stage*, in which a probabilistic surrogate model is used to learn about the objective functions (i.e., the relationship between all possible tACS frequency-phase combinations and the subject’s responses). Typically, non-parametric modelling such as Gaussian process (GP) regression is employed due to its versatility and flexibility (Rasmussen and Williams, 2006). At any given iteration, GP regression using all available observations obtained up to that point is used to predict the objective function across the entire parameter space. Most commonly Bayesian optimization requires that evaluations of the objective function have a scalar response (e.g., (Lorenz et al., 2017a, 2017b, 2016a, 2016b, 2015)). However, for research questions involving human judgement, preferences can be more accurate than ratings as human perception is attuned to evaluate differences rather than absolute magnitudes (Kahneman and Tversky, 1979; Miller, 1956; Seymour and McClure, 2008; Stewart et al., 2001, 2005). To address this problem and exploit the strength of human perception, Brochu et al. (2010) have proposed a Bayesian optimization approach based on discrete preferences. In this approach, participants are presented with two (or more) samples from the parameter space at each iteration, for which they simply indicate their preference. This requires minimal cognitive burden on the participant while more robust results can be obtained than is possible with conventional rating scales (Brochu et al., 2010). By using a Thurstone-Mosteller model with a GP, it is possible to relate binary observations to a continuous function. Figure 3 depicts an example of the 1D parameter space for montage Cz-Oz; it shows how a set of preferences is used at each iteration to infer and update a GP model.

**Figure 3.**
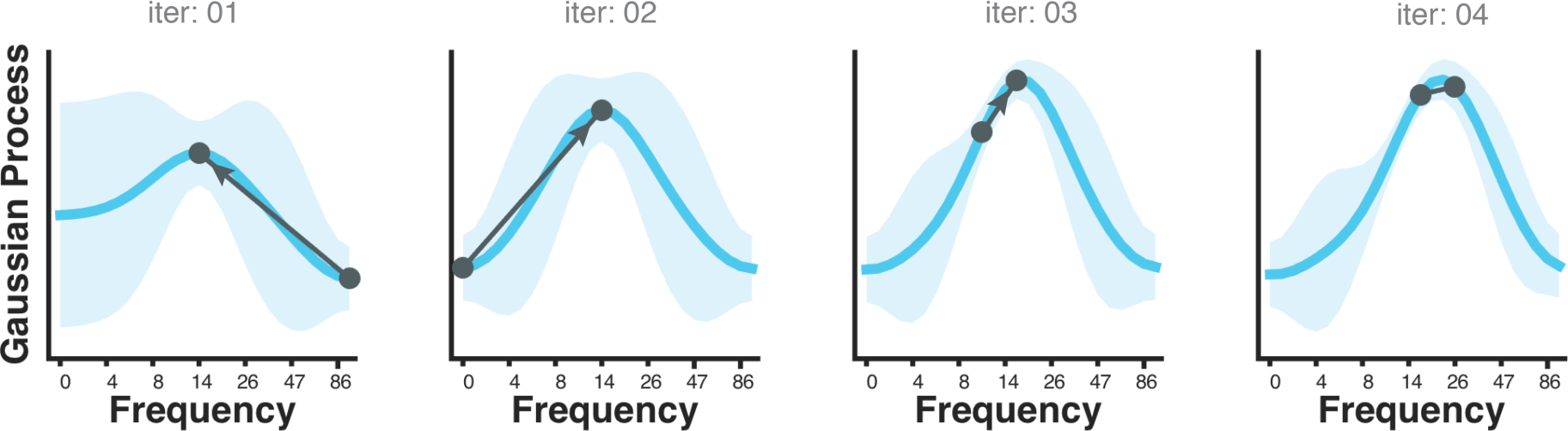
Gaussian process regression based on binary observations. At each iteration (“iter”), subjects indicate their preference between two points from the parameter space (i.e., the direction of the arrow represents preference). This preference serves as the input to a function that models a GP taking into account all preference ratings given up to that point, as well as prior assumptions about the smoothness of the function. It can be seen, that uncertainty (i.e., variance around the mean) about the predictions decreases over time as more preference ratings are available.

The second stage is the *guided search stage*, in which an acquisition function is used to propose a point in the parameter space from where to sample next (i.e., the tACS parameters the subject will receive in the next iteration). This new observation will then be used to update the algorithm’s probabilistic surrogate model. The role of the acquisition function is to guide exploration of the parameter space for achieving its learning goal (i.e., finding the tACS frequency-phase combinations that maximize phosphene intensity) by evaluating the utility of each candidate point (Shahriari et al., 2016). As such, the acquisition function must balance a trade-off between exploring the parameter space and exploiting the current set of parameters for which measurements have already been collected; this allows for an efficient and reliable search over an exhaustive parameter space.

#### Discrete preference GP regression

In order to estimate a GP model relating to preference data, we followed an approach proposed by Brochu et al. (2010). From a methodological perspective, this approach treats preferential data provided by subjects as a classification task. A GP is employed to model the underlying objective function associated with each input. This surrogate model serves to capture the preference for each input such that if input i_1_ is preferred to input i_2_ the surrogate function at i_1_ should be greater than at i_2_. In the study presented here, the distinct inputs correspond to two distinct tACS frequency-phase combinations.

Formally, for a candidate pair of inputs, i_l_ and i_2_, the associated surrogate function is modelled as follows:

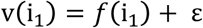

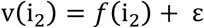

where GP prior to *f*( ) is assigned. Following Brochu et al. (Brochu et al., 2010), the difference in the surrogate function *f*(i_1_) − *f*(i_2_) is studied in order to predict whether input i_1_ will be preferred. Formally, a probit link function is employed in order to obtain a probability that i_1_ is preferred to i_2_ as follows:

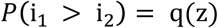

where q( ) is the cumulative distribution function and z is defined as 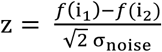. Given a set of training data, fitting such a model corresponds to fitting a GP for a classification task (Bishop, 2007). Due to the non-linear nature of the probit link function an analytic solution is not available for the surrogate function. While a variety of methods have been proposed, here a Laplace approximation was employed as proposed by Brochu et al. (2010); for further details see Bishop (2007). The idea of employing a surrogate function to model binary observations has been employed extensively within the statistics and machine learning literature. When a probit link function is used, as in the present work, such an approach is referred to as Thurstone-Mosteller model.

#### Covariance kernel

GPs are fully specified by their mean and covariance functions. As a prior, we employed a zero mean function and as covariance function we chose an anisotropic squared exponential (SE) kernel (Rasmussen and Williams, 2006). The SE kernel encodes the basic prior assumption that points close in the task space elicit similar responses while points far from each other could exhibit distinct responses:

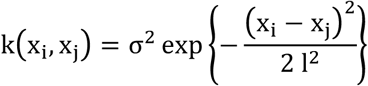

where x_i_ and x_j_ each correspond to an tACS frequency-phase combination, and (x_i_ − x_j_)^2^ determines their Euclidean distance in the parameter space. The parameters σ^2^ and l each determine the variance and length scale of the covariance kernel, respectively; they are referred to as *hyper-parameters* of the covariance kernel. For Study 1, length-scale parameters of the SE kernel were tuned based on a prior offline study, assessing preference ratings across a smaller parameter space (10 frequencies x 6 phases) with eleven participants for montage F4-P4, and five participants for montage O1-O2. Based on the data from Study 1, length-scale parameters were retuned for Study 2. In both cases, hyper-parameters were selected using type II maximum likelihood (Rasmussen and Williams, 2006), implemented via a grid search algorithm (see section “Phase-dependency of phosphene perception” for details). While length-scale parameters differed across studies, the same length-scale parameters were employed for different montages within each study, and were kept fixed for all subjects.

#### Acquisition function

| In the context of preferential data the acquisition function must effectively select two points so that a comparison is possible. Brochu et al. (2010) describe an approach where the first point proposed always corresponds to the current maximum *f*(x^+^), while the second point is then selected by maximizing the expected improvement. The expected improvement (EI) is defined as:

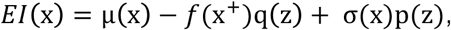

where p( ) is the probability density function, q( ) is the cumulative distribution function and z is defined as: 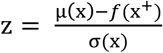, for which μ(x) and σ(x) are the mean and standard deviation of the Gaussian predictive posterior distribution at point x. Since the underlying surrogate function follows a GP, this type of acquisition function can be used to select new candidate points in the parameter space. Informally, this choice of acquisition can be seen as trying to maximize the expected improvement over the current best. Brochu et al. (2010) ran an extensive empirical study and found that the EI acquisition function was preferable to a random acquisition or an acquisition based the point of highest uncertainty.

The initial pair of tACS parameters proposed by the acquisition function was identical across subjects as a set of points was chosen that maximized variance given the prior (Brochu et al., 2010). For proposing points in the parameter space, the acquisition function returned fractions instead of integers; therefore fractions were rounded to obtain desired integer points, corresponding to a pre-defined tACS frequency-phase combination from the parameter space. However, in some rare cases this procedure resulted in the acquisition function proposing the exact same tACS frequency-phase combination for comparison at a given iteration (especially towards end of the run for Study 2, when the acquisition function gained more and more certainty about the optimum). As this was unintended by the experimenters, those iterations were removed from post-hoc analyses; in total this resulted in removing 6 out of 400 iterations in Study 1 (F4P4: 5/200; O1-O2: 1/200), and 47 out of 400 iterations from Study 2 (F4P4: 31/200; O1-O2: 16/200). The removal of these iterations was motivated by the fact that corresponding preference ratings were random and therefore introduced undesirable variance into the GP models.

### Preference probability across parameter space

For an initial analysis, the preference probability for each possible tACS frequency-phase combination was computed across all subjects for each montage and each study separately. For this analysis, the ratio was determined between the number of times a certain tACS parameter combination was preferred over the total number of occurrences; this resulted in a value between 0 (never preferred) and 1 (always preferred) for each point in the parameter space. However, these preference probability maps were not suited for assessing frequency-and/or phase-specific effects in phosphene perception, as they did not account for distances among preference ratings. For example, it was possible that a certain tACS combination (e.g., 19 Hz with 0°) had been more frequently compared to a distant tACS combination (e.g., 12 Hz with 210°), which could have contributed to a high preference probability value because such comparisons are perceptually easier to differentiate than tACS combinations in close proximity (e.g., 19 Hz with 60°) that are more ambivalent. Equally, such a comparison makes it difficult to elucidate whether differences in preference probability can be attributed to frequency, phase, or possibly a combination of both.

Therefore, to better assess frequency- and/or phase-specific effects in phosphene perception, in a subsequent restricted analysis, only a subset of preference probabilities were compared. Here, for assessing frequency-specific effects, only preference ratings were considered that directly compared frequencies with each other that had a relatively similar phase (between 270 and 120° in Study 1 and between 300-60° in Study 2). The intuition behind this approach was, that for these comparisons, the difference in preference probability could be more directly attributed to the difference in frequency than phase (especially for the narrower phase range in Study 2). Equally, for assessing phase-specific effects, only preference ratings were considered that directly compared phases with each other that had a relatively similar frequency (between 14-26 Hz in Study 1, and between 16-22 Hz in Study 2); again this allowed differences in preference ratings to be attributed to the difference in phase instead of frequency. The particular range was chosen heuristically for each study and incorporated a trade-off between the number of preference ratings available and a narrow frequency or phase range.

For statistical inference on these “zoomed-in” preference probabilities, non-parametric permutation testing was performed (10,000 permutations). Preference ratings were randomly shuffled (e.g., while originally a participant indicated 14 Hz with 0° to elicit a stronger phosphene perception than 14 Hz with 180°, this preference might be reversed for certain permutations). Based on these shuffled preference ratings, preference probabilities were computed and stored at each permutation. This was done for each possible frequency (when assessing frequency-specific effects) and phase (when assessing phase-specific effects) and allowed statistical inference about whether obtained preference ratings were significantly below or above the chance level of 0.5. For this analysis, the *α*-level was set to 0.1 because in light of the research question it was equally important to identify frequencies and phases for which there was no detectable difference in phosphene perception (i.e., for which participants were “performing” at chance level ~ 0.5). Therefore, in the present case, an effort was made to decrease the Type I error (“False Negatives”, i.e., to conclude that participants perform below or above chance level, when in fact they are not) by adjusting the Type II error (i.e., *α*-level) slightly up to 0.1 (“False Positives”, i.e., to conclude that participants are performing at chancel level, when in fact they are not). To account for multiple comparisons (as this analysis was done separately for 26/16 different frequencies in Study 1/Study 2 and 12 phases), false discovery rate (FDR) correction (Benjamini and Hochberg, 1995) was performed.

### Group-level Bayesian models

Group-level Bayesian models were obtained for each study separately by inferring the GP based on all available preference ratings of all subjects for a given montage. This analysis was more sensitive for assessing phase-dependent effects than simply studying preference probabilities, because inferred GPs automatically incorporate distance information among preference pairs (see Figure 3). However, the inferred GPs are dependent on the choice of hyper-parameters; therefore, length-scale parameters of the covariance function were selected according to the procedure described below (“Phase-dependency of phosphene perception”). To assess statistical significance of each group-level Bayesian model, one-tailed non-parametric permutation testing was performed. For each of the 1,000 permutations, preference ratings from all subjects were randomly shuffled. Based on these shuffled preference ratings, for each permutation, a new group-level GP was computed and the maximum value of the GP (“height of GP”) was stored. The intuition behind this approach was that the “height of GP” can be directly interpreted as the correspondence of preference ratings across subjects, i.e., the more consistently subjects prefer 19 Hz over 14 Hz, the larger the gradient, thus directly influencing the height of the GP. Based on the distribution of “height” values, p-values were obtained for each group-level Bayesian model.

### Phase-dependency of phosphene perception

As mentioned above, an anisotropic covariance kernel was employed in both studies with two independent kernel length-scale parameters for the two dimensions “Frequency” and “Phase”. To assess if the data from the two studies indicated an effect of phase on phosphene perception, the type II maximum likelihood of the group-level Bayesian models was computed as a function of varying length-scale parameter for “Phase” (i.e. 1,2,3,4,5,7,10,20, and 50), while keeping the length-scale parameter for “Frequency” fixed. The type II maximum likelihood estimation corresponds to a computationally feasible approximation where the log marginal likelihood is maximized with respect to the hyper-parameters (Rasmussen and Williams, 2006):

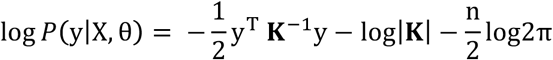

From this equation, it can be seen that the log marginal likelihood is conditioned on the parameters of the kernel θ. The equation consist of three easily interpretable terms:

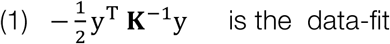

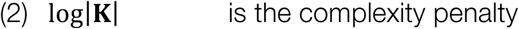

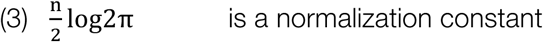

While the data fit decreases monotonically with the length-scale as the GP model become less flexible, the penalty term decreases with the length-scale as the GP model becomes less complex. Therefore, the log marginal likelihood automatically incorporates a trade-off between model fit and model complexity (as it takes the data fit minus the penalty term), and higher values are desirable.

Directly comparing log marginal likelihood for varied “Phase” length-scale parameters, thus serves as a good indicator for assessing phase-dependent effects of phosphene perception. While high log marginal likelihood values for small to medium length-scales (i.e., 1-10) would indicate a phase effect, high log marginal likelihood values for large length-scale parameters (i.e., 20-50) would contradict a phase-dependent effect, as such smooth kernels would result in similar predictions along the “Phase”-axis.

### Difference between montages

To assess if phosphene perception revealed statistically different optimal tACS parameters between montages F4-P4 and O1-O2, non-parametric permutation testing was performed. For this purpose, the Euclidean distance (ED) was computed between the predicted optimum of the two montages (which was 1, see Results). To obtain an appropriate null distribution, group-level Bayesian models were re-calculated by permuting the montage “labels” of subjects, and ED measures between montages were stored for each permutation; the number of possible permutations was 2^10^ (as *n*=10), Based on this generated distribution, a p-value was obtained for the empirically obtained ED value.

### Volume conductor model

Current density on the retinas for each electrode montage was approximated computationally using an anatomically realistic volume conductor model of a 34 year-old male (Christ et al., 2010). The details of the computational procedure have been reported previously in Laakso and Hirata (2013). Briefly, the electric scalar potential was determined using the finite-element method with first-order cubical elements (0.5 mm side length), and the current density was calculated from the gradient of the scalar potential. The electrodes were modelled as rubber pads (0.1 S/m), the size and thickness of which were identical to the electrodes used in the experiments. A 2.5 mm thick layer of conductive electrode paste (1.6 S/m) was added between the electrodes and the scalp. Return electrodes were modelled using the Neumann boundary condition in the neck.

The effect of phase was modelled by running two computer simulations for each electrode montage. In each simulation, only one pair of electrodes was active and the other pair was modelled as passive conductors. The total current for an arbitrary phase difference was calculated as a linear combination of the currents of the electrode pairs. This approach is valid because the currents are relatively weak and thus tissue electrical conductivities can be assumed to be linear.

## Results

### Replication of frequency-dependent phosphene perception

In Study 1, montage Cz-Oz was tested using a 1D parameter space to validate the optimization technique against previous studies that employed the same montage to assess phosphene perception with conventional rating scales (e.g., (Kanai et al., 2008)). The probability of preference map across the frequency space pooled across all subjects is depicted in Figure 4a. It was observed that the acquisition function predominantly sampled frequencies in the alpha and beta ranges, i.e., 9-36 Hz while omitting frequencies in the delta (1-3 Hz), theta (4-7 Hz), or gamma ranges (> 40 Hz). The efficiency of the algorithm’s sampling behaviour reflects the smoothness assumptions encoded in the GP kernel: as the corners of the parameter space (i.e., 0 Hz and 100 Hz) were assessed by the algorithm and subjects did not indicate any preference for these, the GP model predicted low values for frequencies encompassed in the delta, theta, or gamma ranges; this in turn affected the guided search to focus on more “optimal” frequencies in the alpha and beta ranges for inducing strong phosphene perception.

**Figure 4.**
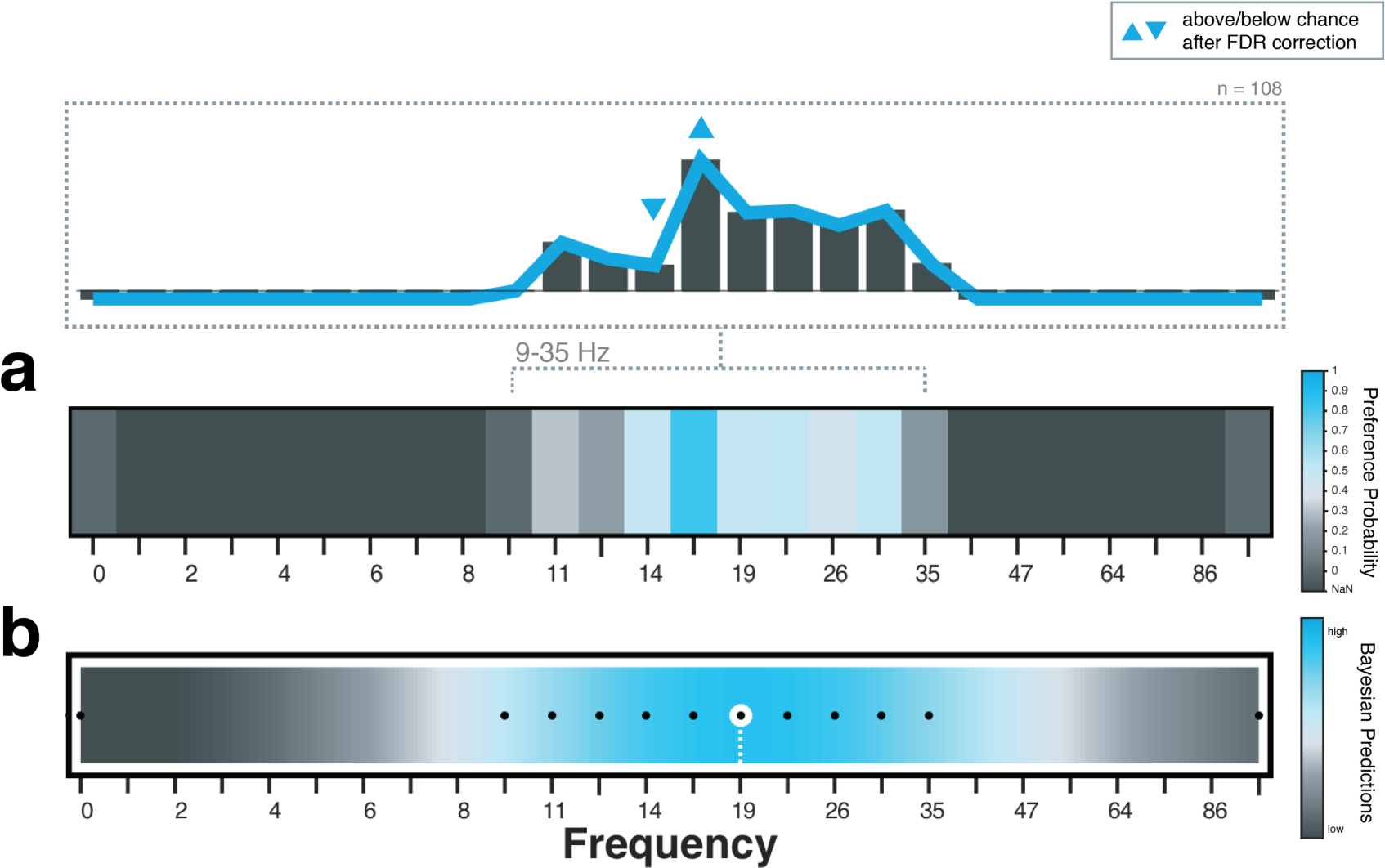
Group-level results for montage Cz-Oz. (a) Preference probability across 1D frequency space. Blue indicates higher preference probability; dark grey corresponds to points not sampled by the acquisition function (NaN). The bar plot on top indicates zoomed-in preference probabilities when only taking into account comparison in the range between 9-35 Hz. Blue triangles indicate statistical significance, while “negative” bars correspond to NaN (i.e., no direct comparisons available). (b) Group-level Bayesian model across 1D frequency space. Blue indicates Bayesian predictions of higher perceived phosphene intensity. Black dots correspond to points sampled by the acquisition function and the white dashed line indicates the tACS frequency with the highest perceived phosphene intensity.

When “zooming” into the preference ratings and just taking into account frequencies between 9 and 35 Hz, phosphenes at 14 Hz were rated significantly below chance level while phosphenes at 16 Hz were rated significantly above chance level (bar plot in Figure 4a). This is consistent with Kanai et al. (2008), which demonstrated a steep fall in phosphene intensity ratings between 14 and 16 Hz. The group-level Bayesian predictions across the parameter space are depicted in Figure 4b. The strongest phosphene perception was predicted at 19 Hz (the maximum predicted value is marked as a white dashed line). Again, this finding is in line with previous literature reporting the strongest phosphene perception at 19 Hz (Kanai et al. (2008)). The successful replication of frequency-dependent phosphene perception validates the Bayesian optimization approach based on preference ratings for assessing tACS-induced phosphene perception.

### Phase-dependent phosphene perception

The probability of preference across the parameter space pooled across all subjects for montage F4-P4 and O1-O2 is depicted in Figure 5. For Study 1 (Figure 5a), the acquisition function sampled densely around the predicted optimal tACS frequency range in the upper alpha and lower beta band (12-19 Hz) with little to no exploration of frequencies in the delta, theta, lower alpha (< 11 Hz), higher beta (> 19 Hz) or gamma ranges. This indicated a clear effect of frequency in phosphene perception for these two montages; with higher preference ratings for frequencies between 14 and 19 Hz. In addition, phases between 300° and 90° were more densely sampled. Although the edges along the “Phase”-axis, i.e., 180° and 210° have also been sampled repeatedly, those were less preferred by the subjects. When taking a closer look into the preference ratings and only taking into account the frequency range from 14-26 Hz, a clear phase effect was found, with preference ratings significantly below chance level for 180° and 210° for both montages. By contrast, for 30° relative phase, preference ratings were found to be significantly above chance. Results of Study 1 pointed towards a very exploitative behaviour of the acquisition function, presumably driven by (too) large length-scale parameters of the covariance kernel.

**Figure 5.**
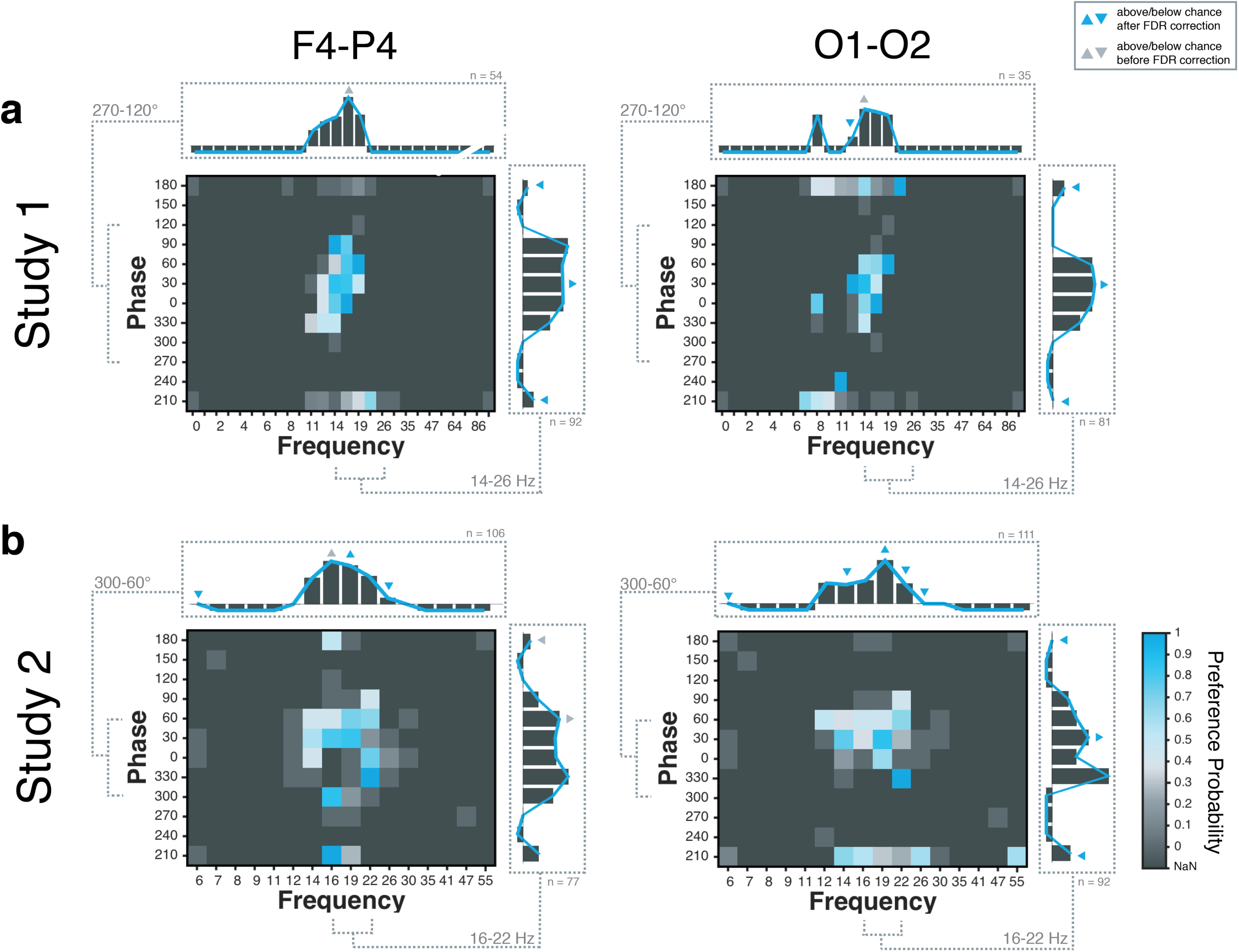
Group-level preference probability for montages F4-P4 and O1-O2. Preference probability and zoomed-in preference probability (bar plots) illustrating frequency-dependent (top) and phase-dependent-effects (right side) of phosphene perception for (a) Study 1 and (b) Study 2. Blue indicates higher preference probability; dark grey corresponds to points not sampled by the acquisition function (NaN). For the bar plots, blue triangles indicate statistical significance, while “negative” bars correspond to NaN (i.e., no direct comparisons available).

Study 2 was conducted to build on work from Study 1, and allow for more exploration and more precise sampling along the frequency and phase axes. For Study 2, the hyper-parameters were retuned (based on the results of Study 1), resulting in smaller length-scale parameters, explaining the more explorative behaviour of the acquisition function for Study 2 (Figure 5b). In line with Study 1, for both montages a frequency-dependent effect was found, with higher preference values for phosphenes at frequencies between 14-22 Hz. A steep fall in phosphene intensity was observed between 22 Hz and 26 Hz in both montages. When considering frequencies ranging from 16-22 Hz, a clear phase effect was found for the O1-O2 montage, with preference ratings significantly below chance for 180° and 210°. This phase-effect was less pronounced for the F4-P4 montage.

In sum, while these results point towards a phase-dependent effect of phosphene perception, it should be noted that this analysis did not take into account the spatial relationship between pairs of tACS parameters (i.e., that 19 Hz-0° compared to 19 Hz-60° is far closer in the parameter space than 9 Hz-0° compared to 22 Hz-60° and should have more influence for studying phase-dependent effects). Therefore, in a subsequent analysis group-level Bayesian models were obtained based on all available preference ratings per montage. The log marginal likelihood of these group-level models was computed as a function of varying length-scale parameter for “Phase”, while keeping the length-scale for “Frequency” fixed. Results of these analyses are listed in Table 1 and indicate a clear effect of phase for both montages as medium-sized length-scale parameters (i.e., 3 for Study 1 and 7 for Study 2) resulted in higher log marginal likelihood values than large length-scale parameters (i.e., 20 or 50). The slightly smaller length-scales found for Study 1 could be explained by the small number of comparisons available across the “Phase”-dimension, due to the more exploitative behaviour of the acquisition function in Study 1.

**Table 1.** Log marginal likelihood values for different length-scale parameters (1-50) of kernel for the “Phase” dimension (underlined values correspond to maximum log marginal likelihood values for each study)

|  | 1 | 2 | 3 | 4 | 5 | 7 | 10 | 20 | 50 |
| --- | --- | --- | --- | --- | --- | --- | --- | --- | --- |
| Montage F4-P4 |  |  |  |  |  |  |  |  |  |
| Study 1 | -3.95 | -0.02 | <u>0.64</u> | -1.55 | -3.75 | -9.65 | -33.40 | -93.40 | -127.88 |
| Study 2 | -50.89 | -52.06 | -37.36 | -25.92 | -20.87 | <u>-20.69</u> | -24.00 | -29.49 | -31.71 |
| Montage O1-O2 |  |  |  |  |  |  |  |  |  |
| Study 1 | -18.40 | -9.93 | <u>-6.03</u> | -7.19 | -11.15 | -25.32 | -53.55 | 106.34 | 135.21 |
| Study 2 | -17.87 | -31.62 | -27.02 | -19.39 | -15.85 | <u>-14.49</u> | -15.20 | -16.86 | -17.65 |

### No detectable differences between montages

Group-level Bayesian models for each montage are depicted in Figure 6; the length-scale parameters of “Phase” were selected based on the results listed in Table 1 (i.e., 3 for Study 1 and 7 for Study 2) and the length-scale parameter of “Frequency” was identical across studies and montages. Using permutation testing, statistical inference of each group-level Bayesian model was obtained based on the maximum predicted value (“height” of the GP). This analysis revealed significant group-level models for each montage and each study (p < .001), indicating strong inter-subject consistency in the preference ratings.

**Figure 6.**
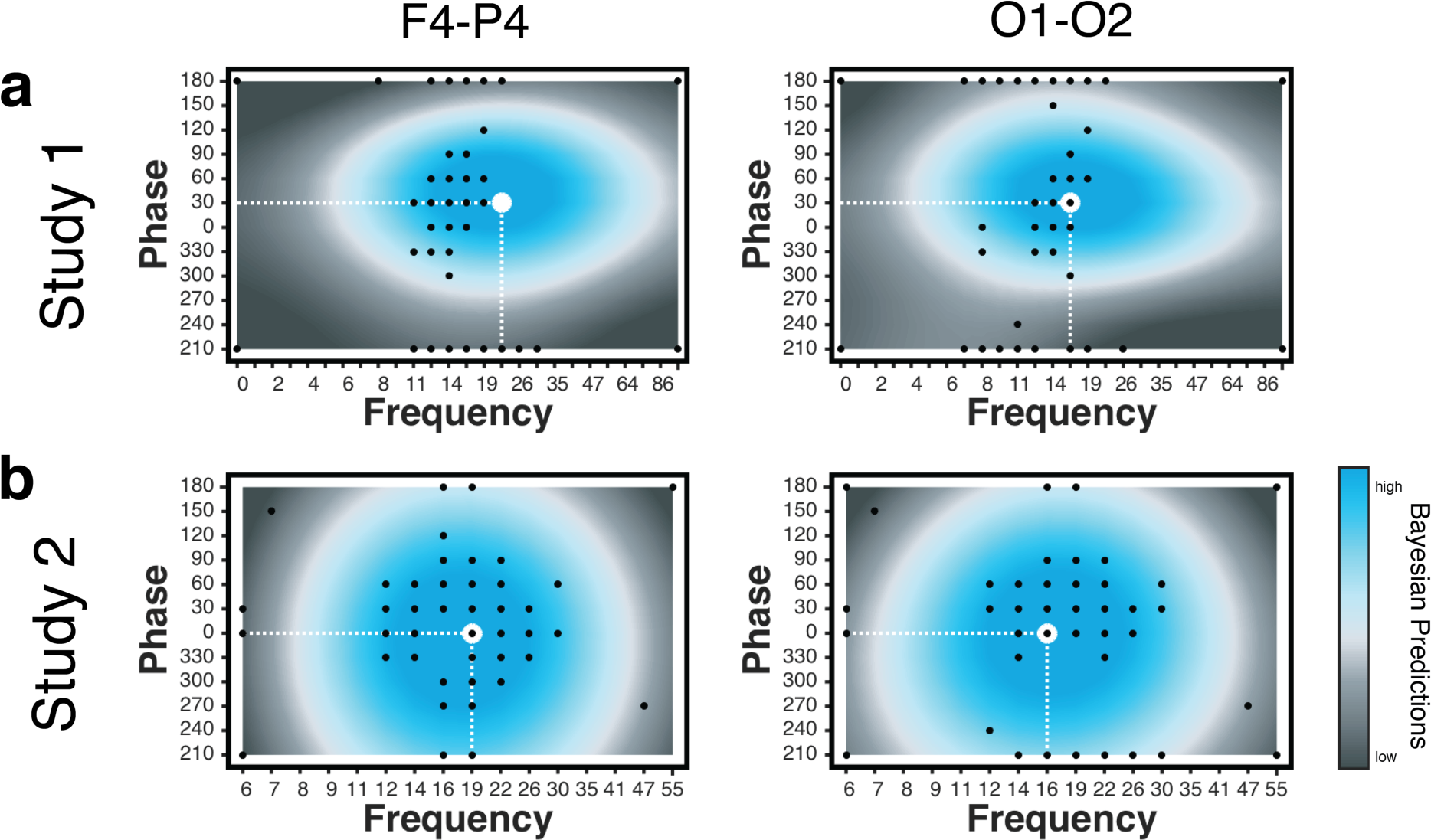
Group-level Bayesian model for montages F4-P4 and O1-O2. Group-level results indicate phase-dependent effects of phosphene perception in (a) Study 1 and (b) Study 2. Blue indicates higher perceived phosphene intensity. Black dots correspond to points sampled by the acquisition function and the white dashed line indicates the tACS frequency-phase combination with the highest perceived phosphene intensity.

The tACS frequency-phase combination associated with the strongest phosphene perception (i.e., the maximum predicted value) is marked as a white dashed line in Figure 6. For montage F4-P4, the optimum was predicted at 22 Hz with 30° for Study 1 and at 19 Hz with 0° for Study 2. For montage O1-O2, both studies predicted the optimum at 16 Hz, either with 30° phase difference (Study 1) or 0° (Study 2).

Besides a small difference in the optimal predicted frequency between montages, visual inspection of the Bayesian models in Study 2 does not indicate any difference in phase-dependency of phosphene perception between the two montages: for both F4-P4 and O1-O2, phases between 210-240° and 120-180° are predicted to induce lower phosphene perception than phases between 270-60°. To assess if the difference in optimal frequency (16 Hz for O1-O2 vs. 19 Hz for F4-P4) was statistically significant for Study 2, non-parametric permutation testing was performed. This analysis revealed no significant difference between the two montages (*p* = .96).

### Empirical results are validated by computational model

The empirical data obtained were subsequently compared to the results of a computational volume conductor model predicting current density in the retinas based on anatomical information of electrode placement (Laakso and Hirata, 2013) as well as differences in relative phase. Results of the computational analysis are depicted in Figure 7. We found that currents near the eye are in the superior-inferior direction for all montages, and that the highest current densities are found in the upper and lower peripheries of the retinas (Figure 7a-c). This is in line with the subjective reports of our participants in both studies, who exclusively reported phosphenes in their visual periphery (see Supplementary Figure 1 for drawings of participants visualizing the extent of their phosphene perception). Both F4-P4 and O1-O2 montages yield the highest retinal current density when tACS stimulation is applied in phase (i.e., 0°), and the lowest current density when the stimulation is applied at opposite phase (i.e., 180°). The computational results validate our empirical data, highlighting a phase-dependent effect of phosphene perception. When comparing across different montages (Figure 7d), we found that the F4-P4 montage produced higher retinal current densities than the O1-O2 montage. The Cz-Oz montage produced weaker retinal currents than both the F4-P4 and O1-O2 montages.

**Figure 7.**
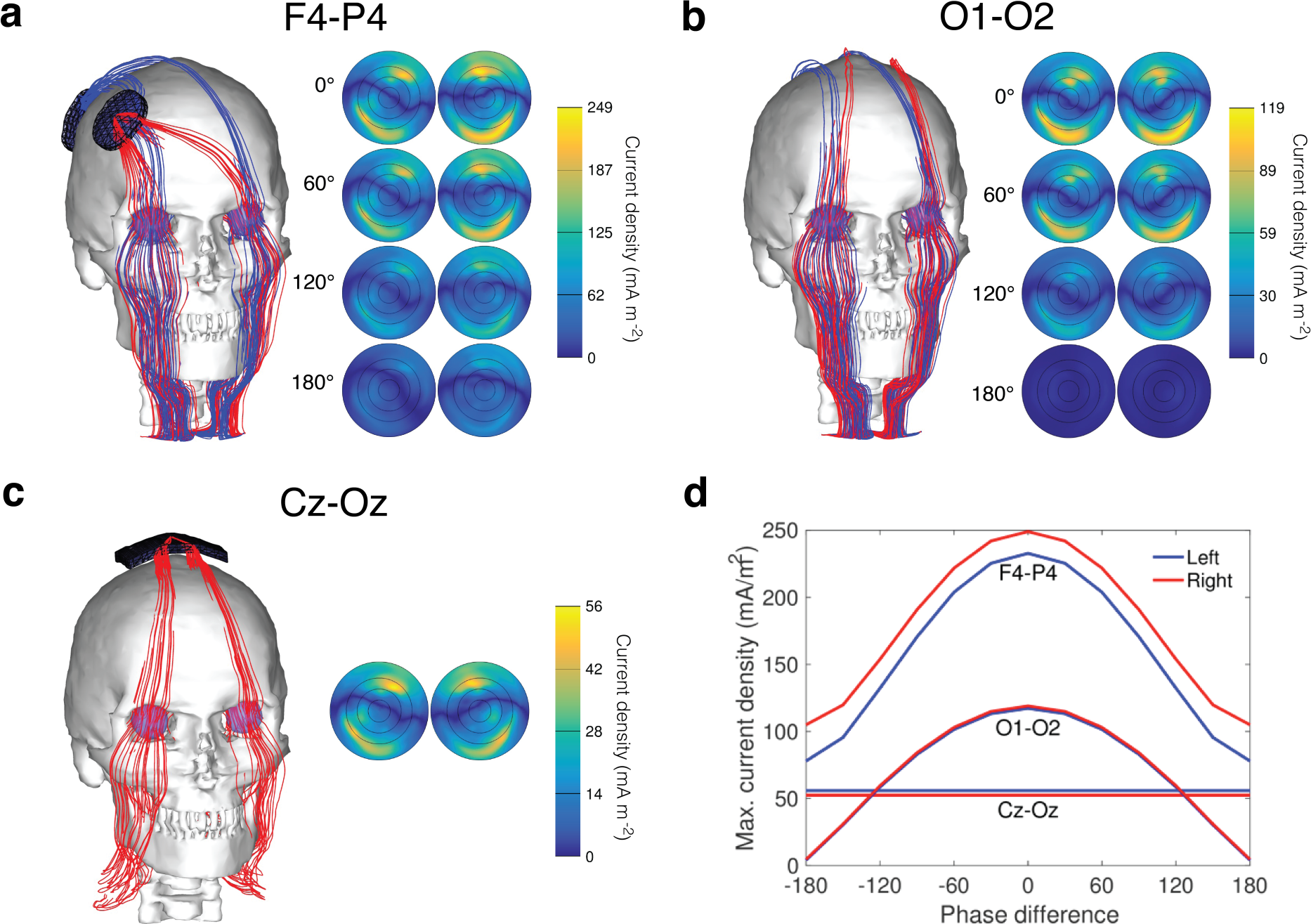
Computational modeling results for different montages. (a-c) Left panel: Streamline visualization showing the direction of current for montage. Only streamlines passing through the eyes are shown. Red and blue streamlines show the currents originating from the two stimulation electrodes, i.e., (a) F4-P4 and (b) O1-O2. Right panel: Current distribution on left and right retinas (posterior view) for four different phase differences (0°, 60°, 120° and 180°). The component of the current density vector perpendicular to the retina is shown. (d) Comparing maximum retinal current density across montages and for different phases.

## Discussion

In this study we demonstrated that Bayesian optimization based on binary preference ratings provides a feasible and efficient method for assessing tACS-induced phosphene perception across a large tACS parameter space (312 possible combinations in Study 1 and 192 in Study 2). While hypothetically testing each possible tACS parameter combination would have taken up to 83 min in Study 1 and 51 minutes in Study 2, the proposed method took 5 minutes per participant.

In Study 1, results obtained with montage Cz-Oz could be directly compared with previous studies employing the same montage and assessing phosphene perception using conventional rating scales. The optimal frequency of 19 Hz identified by the optimization procedure was highly consistent with results of Kanai et al. (2008), serving as an excellent demonstration of the validity and reliability of the method. Frequency-dependent effects on phosphene perception were also found for montages F4-P4 and O1-O2 in both studies. This finding was expected, as studies employing different montages (e.g., left motor cortex-right orbitofrontal cortex (Turi et al., 2013)) have also reported that stimulation frequencies close to 19 Hz induce the strongest phosphene perception. Frequency-dependent effects of phosphene perception can be understood from response properties of retinal ganglion cells, which are known to have a temporal frequency tuning with a peak sensitivity in the beta frequency band in light conditions that shifts towards the alpha-range in the dark (Kaplan and Benardete, 2001; Kar and Krekelberg, 2012).

Besides replicating frequency-dependent effects, this study is the first to assess phase-dependent effects in tACS phosphene perception for two different montages (i.e., F4-P4 and O1-Oz). We found that phosphenes were perceived more intensely for relative phase differences between the two stimulation electrodes closer to 0° (270-60°) and less intensely when the relative phases were more temporally lagged (90-240°). These empirical observations were validated by a computational volume conductor model showing that the phase-specific effect of phosphene perception could be explained by a reduced current density on the retinas for less synchronized oscillations. This phenomenon is known as interference in wave propagation, which describes the superposition of more synchronized oscillations resulting in a wave of higher amplitude (constructive interference) while oscillations that are out of phase form waves of lower amplitudes (deconstructive interference).

Our results indicate that the effect of phase differences on phosphene perception is a relatively subtle perceptual phenomenon, less pronounced than frequency-dependent effects; therefore, more conventional methods, such as rating scales may have lacked sufficient sensitivity to assess these relatively subtle differences in human perception. Moreover, our precise computational model taking into account both the anatomic location of electrode placement as well as differences in phase could provide a useful tool for guiding electrode placement for future tACS studies.

The finding of phase-dependent phosphene perception has important implications for future studies drawing conclusions about phase-related effects on cognitive or behavioural performance within the “phosphene-critical” frequency range, i.e., frequencies encompassing the alpha and beta band that have been shown to elicit strong phosphene perception. This is a timely topic, with many studies focusing on phase-related rather than frequency-related effects of tACS on cognitive and behavioural performance (Guerra et al., 2016; Helfrich et al., 2014; Polanía et al., 2012, 2015; Violante et al., 2017). Alongside concerns about the possible entrainment effect induced by retinal phosphenes (Schutter, 2016), the perception of phosphenes may also modify the alertness of participants, and thus could possibly alter attentional processes (Turi et al., 2013). Consequently, any cognitive or behavioural effects of relative phase found for frequencies in the alpha or beta band, may be potentially be driven by changes in the perceived phosphene intensity rather than desired effects of neural entrainment. This highlights the need for developing appropriate control conditions when interested in studying phase-related effects of tACS. While to date photic stimulation has been proposed (Mehta et al., 2015) to disentangle if tACS-induced behavioural effects are due to retinal or neural entrainment over the stimulation site, photic stimulation may not be sensitive enough to properly control for the subtle perceptual differences associated with different phases. Adding to the complexity, the intensity of phosphene perception varies across montages and may also interact with changes in the subject’s cognitive load (Cabral-Calderin et al., 2016; Violante et al., 2017; Vosskuhl et al., 2016). Therefore, we advise to characterize the perception of phosphenes on a study-by-study basis, which can be easily and efficiently achieved through Brochu et al.’s (2010) Python implementation of the Bayesian optimization approach available on https://github.com/misterwindupbird/IBO.

Beyond that, our approach could be used to include stimulation intensity as an additional dimension to be searched through, allowing for the rapid detection of individual thresholds of phosphene perception. An interesting avenue for future work would be to use our technique to empirically compare phosphene perception across different montages; however this would require further technological advances in the development of high-density transcranial electrical stimulation caps in order to remotely control and test different combinations of active and reference electrodes.

From a methodological point of view, the less explorative sampling behaviour of the acquisition function in Study 1 highlights the need for appropriate priors when employing Bayesian optimization based on preference ratings. As a rule of thumb, smaller kernel sizes are preferred when more exploration is desired. In addition, measures need to be taken to prevent the acquisition function from proposing the exact same tACS frequency-phase combination for comparison at a given iteration.

In summary, Bayesian optimization has proved to be a valuable tool for assessing subtle perceptual differences in tACS-induced phosphene perception. This approach provides a reliable method to model complex response functions solely based on preference ratings. By intelligently sampling the parameter space and only choosing the most informative comparisons, this approach combines high efficiency with the strength of human perception. While this approach has originally been proposed for facilitating realistic material design in the field of computer graphics (Brochu et al., 2010), this study may pave the ways to a broader application in cognitive science and psychology.

## Acknowledgements

This work was supported by the NIHR Imperial BRC and the Leverhulme Trust. IRV is funded by the Wellcome Trust (103045/Z/13/Z).

## Supplementary Material

**Supplementary Figure 1.**
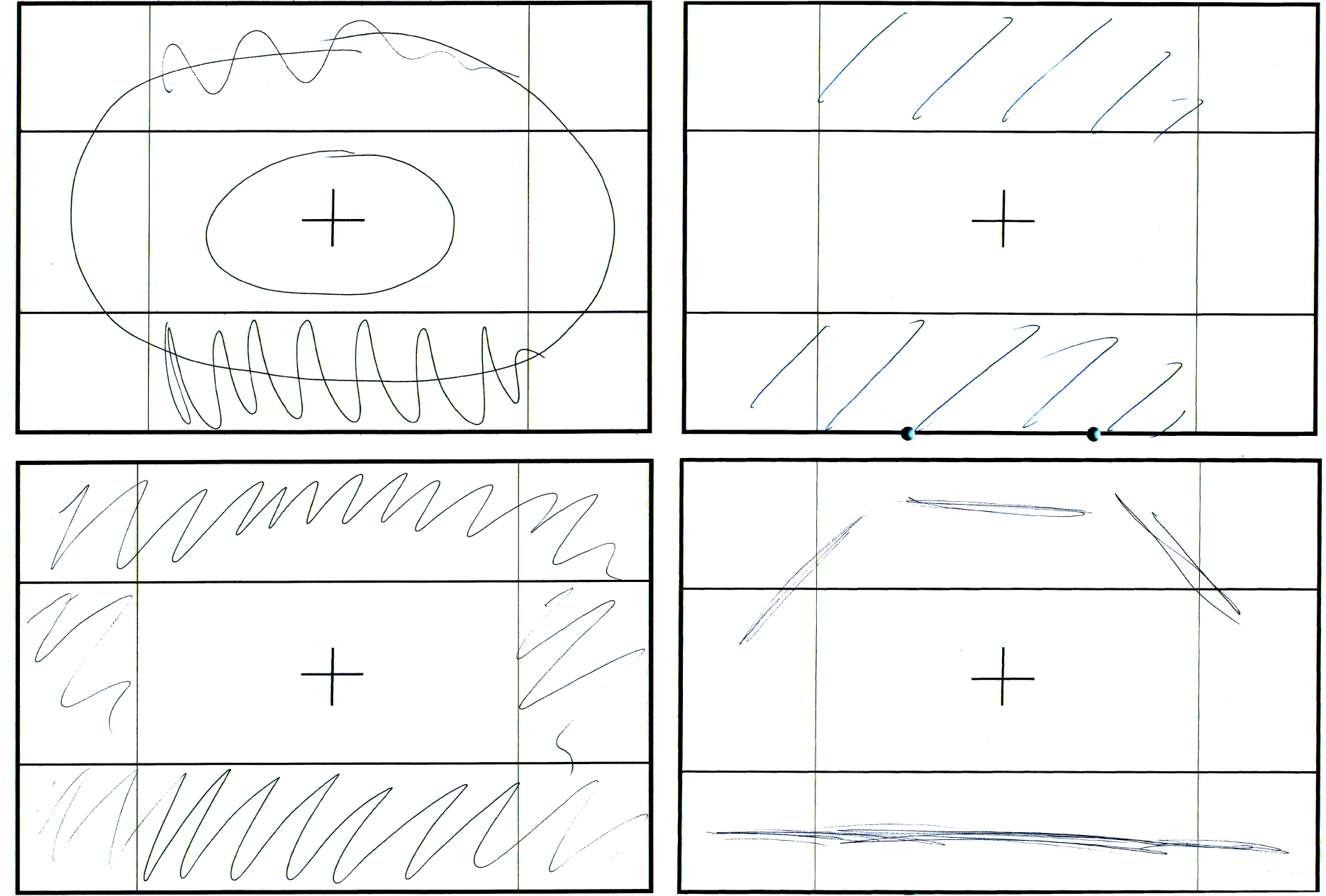
Four examples of participants’ drawing about where in their visual field they had perceived phosphenes. Subjective results were in line with our computational models predicting the highest current density in the periphery of the subjects’ retina.

## Footnotes

1 The standard procedure to assess tACS-induced phosphene perception is to expose participants to the theoretically assumed strongest tACS frequency (such as 16 Hz) before the experiment starts, thereby acting as a “reference” for subsequent rating blocks. However, this requires for subjects to recall the intensity of the reference phosphene intensity while being exposed to blocks of other frequencies. Thus, this procedure is less precise for determining subtle differences in phosphene perception.

